# Sequence and epigenetic characterization of chromosome 21 centromeres in a family with recurrent Trisomy 21

**DOI:** 10.64898/2026.08.31.748295

**Authors:** F. Kumara Mastrorosa, Kendra Hoekzema, Marcelo Ayllon, Periklis Makrythanasis, Stylianos E. Antonarakis, Evan E. Eichler

## Abstract

Trisomy 21 (T21) is the most common genetic cause of intellectual disability, yet the molecular mechanisms underlying maternal meiosis I errors—responsible for ~70% of free T21 cases— remain poorly understood. In this preliminary study, we used long-read sequencing and genome assembly to investigate the DNA sequence and epigenetic features of chromosome 21 (chr21) centromeres in a family with recurrent free T21 due to maternal meiosis I errors. The mother, who had two affected and three unaffected children, showed no mosaicism or structural rearrangements. One of her two chr21 centromeres lacked a pronounced centromere dip region (CDR), displaying instead a diffuse hypomethylation pattern (dCDR) with much higher methylated CpG levels (55%) compared to its homologue (36%). This dCDR was transmitted to an unaffected child and the affected proband, suggesting it was present in one of the maternal chr21 since she was at least 32 years of age. Chr21 dCDRs were not observed in seven young mothers with children with T21 or previously described in the literature in 108 population haplotypes. We hypothesize that dCDRs may weaken kinetochore function, increasing nondisjunction risk, and propose two models linking such epigenetic variation to maternal age-related T21 risk. These findings highlight the value of complete centromere characterization in families with children with T21 and suggest centromere methylation status of chr21 as a potential T21 risk factor for future investigation.

## INTRODUCTION

Trisomy 21 (T21; MIM: 190685) is the chromosomal condition causing Down syndrome, the most common genomic condition in humans causing intellectual disability (1). Free T21, which accounts for 95% of the cases, refers to the presence of three distinct chromosomes 21 (chr21) as opposed to T21 arising from inheritance of a Robertsonian translocation. Approximately 70% of free T21 cases are due to an error in maternal meiosis I (MMIE) (1), and the affected individuals carry two chr21 haplotypes. Other cases are due to errors in MMIIE, paternal meiosis, or parental mosaicism. Despite being described phenotypically (2) and chromosomally (3) several decades ago, the genetic risk factors and mechanisms leading to T21 are still unclear. The most well-known contributing risk factor is maternal age (4) with a direct correlation with T21 pregnancies. Many hypotheses have been proposed to explain this phenomenon, including age-associated reduced chiasma in oocytes (5,6), cohesin deficiency (7), variation in the size of the chr21 centromeric alphoid DNA (8), and others (9). It is also possible that more than one of these genetic factors may, in fact, contribute to the overall risk.

Centromeres have been proposed to play a role in several human trisomy syndromes because they ensure faithful chromosome segregation during mitosis and meiosis. Multiple studies revealed the importance of kinetochore-associated proteins in preserving the centromere function in mice and human cells (10,11). Specifically, it is widely accepted that the histone H3 variant CENP-A is essential for kinetochore function (12–14). Currently, the relationship between these proteins, the centromere, and the extent of methylation is being investigated. Although variable across centromeres, human alphoid arrays are typically highly methylated (~50 to ~70%) and, in most cases (94%), contain one hypomethylated region (ranging from 90 to 630 kbp; average 240 kbp)—called the centromere dip region (CDR)—where the 5-methyl cytosines (5mCs) transition to a 10-30% methylation percentage (15) and CENP-A accumulates. These features can now be accessed with long-read sequencing technologies coupled with the latest genome assembly algorithms, which allow complete sequence resolution of the centromere higher-order repeat (HOR) array of centromeres (16,17) along with precise definition of the hypomethylated CDR. Recently, we demonstrated that these methods can successfully assemble all three chr21 centromeres of individuals with T21 and performed a large-scale chr21 centromere survey in probands and the general population (18). Here, we sequence and epigenetically characterize the chr21 centromeres transmitted in a family with free T21. This pedigree allows us to track the inheritance of maternal centromeres to each child and observe the genetic and epigenetic variation of the alphoid sequences over time. The family studied has two affected individuals with free T21. We hypothesize that it represents a rare example of a family where there is increased risk for T21 and, as such, it may provide clues into the genetic/epigenetic risk factors contributing to the condition.

## MATERIAL AND METHODS

### Sample recruiting

This family was initially studied in 1992 (19) (family RDS07) and recontacted for this study. After genetic counseling and explanation of the study by a medical geneticist, the family members signed an informed consent (provided by the University of Athens St. Sophia Hospital) and donated EDTA blood.

### UL-ONT sequencing

Ultra-high-molecular-weight (uHMW) gDNA was extracted from blood samples. The white blood cells were isolated from the whole blood with multiple washes with RBC lysis buffer (Qiagen; Venlo, Netherlands) and pelleting. The remaining white blood cells were lysed in a buffer containing 10 mM Tris-Cl (pH 8.0), 0.1 M EDTA (pH 8.0), 0.5% w/v SDS, and 20 mg/mL Rnase A for 1 hour at 37°C. 200 ug/mL Proteinase K was added, and the solution was incubated at 50°C for 2 hours. DNA was purified via two rounds of 25:24:1 phenol-chloroform-isoamyl alcohol extraction followed by ethanol precipitation. Precipitated DNA was solubilized in 10 mM Tris (pH 8.0) containing 0.02% Triton X-100 at 4°C for two days. Libraries were constructed using the Ultra-Long DNA Sequencing Kit (SQK-ULK114) (ONT; Oxford, UK) following the manufacturer’s protocol. Next, 75 uL of the library was loaded onto a primed FLO-PRO114M R10.4.1 flow cell for sequencing on the PromethION, with two nuclease washes and reloads after 24 and 48 hours of sequencing. Sequence reads >100 kbp in length were classified as ultra-long. All ONT data were basecalled using Guppy (v6.3.7 or newer) or Dorado (v0.4.2 or newer) with the SUP model, which both produce sequencing and methylation data.

### HiFi sequencing

DNA quantity was assessed with Qubit dsDNA HS (Q32854) (Thermo Fisher; Waltham, MA, USA) on DS-11 FX (Denovix; Wilmington, DE, USA), and size distribution was checked using FEMTO Pulse (M5330AA & FP-1002-0275) (Agilent; Santa Clara, CA, USA) uHMW DNA was sheared with Megaruptor 3 (B06010003 & E07010003) (Diagenode, Liège, Belgium) using setting 28/30. The resulting DNA was used to generate PacBio HiFi libraries via the SMRTbell Prep Kit 3.0 (102-182-700) (PacBio; Menlo Park, CA, USA) Size selection was performed using diluted (35% v/v) AMPure PB Beads with a ratio of 3.1x to progressively remove all DNA fragments below 10 kbp. Samples were sequenced on the Revio platform on SMRT Cells 25M (PacBio, 102-817-900) with Adaptive Loading and 30-hour movies. Coverage targeted a minimum of 30× in PacBio HiFi reads for the parent and siblings, (1-1.5 SMRT Cells per sample) and 60× for the proband (2-2.5 SMRT Cells, assuming a genome size of 3.1 Gbp).

### Genome assembly and centromere analysis

HiFi and UL-ONT data were both used to assemble the genome of each individual using hifiasm 0.19.9 (20). Mother II-2 chr21 centromeres were identified aligning their assembly to CHM13 with minimap2 (21) v2.24 and following commands: minimap2 -t 8 -I 15G -a --eqx -x asm20 -s 5000. Contigs containing chr21 centromeres mapped to the CHM13 chr21 centromere and extended to the q-arm. CenMAP (v0.3.1) (https://github.com/logsdon-lab/CenMAP), an automated workflow for centromere characterization, was used to characterize the centromeres in all samples. In short, RepeatMasker (22) was used to study the coordinates of the alpha satellite arrays and HumAS-HMMER (https://github.com/fedorrik/HumAS-HMMER_for_AnVIL) was used to characterize the HOR monomers of each centromere. HiFi reads were aligned back to their respective assembly using pbmm2 v1.14.99 (https://github.com/PacificBiosciences/pbmm2) with the following command: pbmm2 align --log-level DEBUG --preset SUBREAD --min-length 5000 --strip -j 4. NucFlag (https://github.com/logsdon-lab/NucFlag) was used to identify assembly errors from these alignments.

Centromere alpha satellite sequences were extracted using seqtk v1.4 (https://github.com/lh3/seqtk) and aligned with minimap2 (21) with following command: minimap2 -x asm20 -c --eqx -secondary=no. The resulting PAF files were mined to extract identity percentage.

CDR-Finder (23) was used to investigate the read coverage, the centromere methylation profiles, and characterize the CDRs. CDR boundaries were refined, visualizing the alignment via Integrative Genome Browser (24). Methylation levels were quantified by calculating the percentage of methylated CpGs–CpGs for which 70% of sequencing reads showed a signature of methylation within each haplotype CDR window. For haplotypes that have multiple CDR windows, only the number of CpGs inside each dip was considered. The cumulative CDR size was calculated adding the size of each CDR.

## RESULTS

To investigate chr21 centromere transmission over time, we recruited a family with five offspring and recurrent T21. Three unaffected children were born at maternal ages 27, 32, and 35, while two affected siblings (a male and a female) were delivered at maternal ages 33 and 38. Blood samples were available for the unaffected mother (I-2), the first two unaffected siblings (II-1 and II-2), and the latest proband (II-5) (**Fig. 1A**). We extracted the DNA from these samples for both PacBio HiFi and ultra-long (UL) Oxford Nanopore Technologies (ONT) sequencing (**Table S1**). These sequencing data were used to independently assemble the genome of each individual using hifiasm v0.19.9(20).

**Figure 1.**
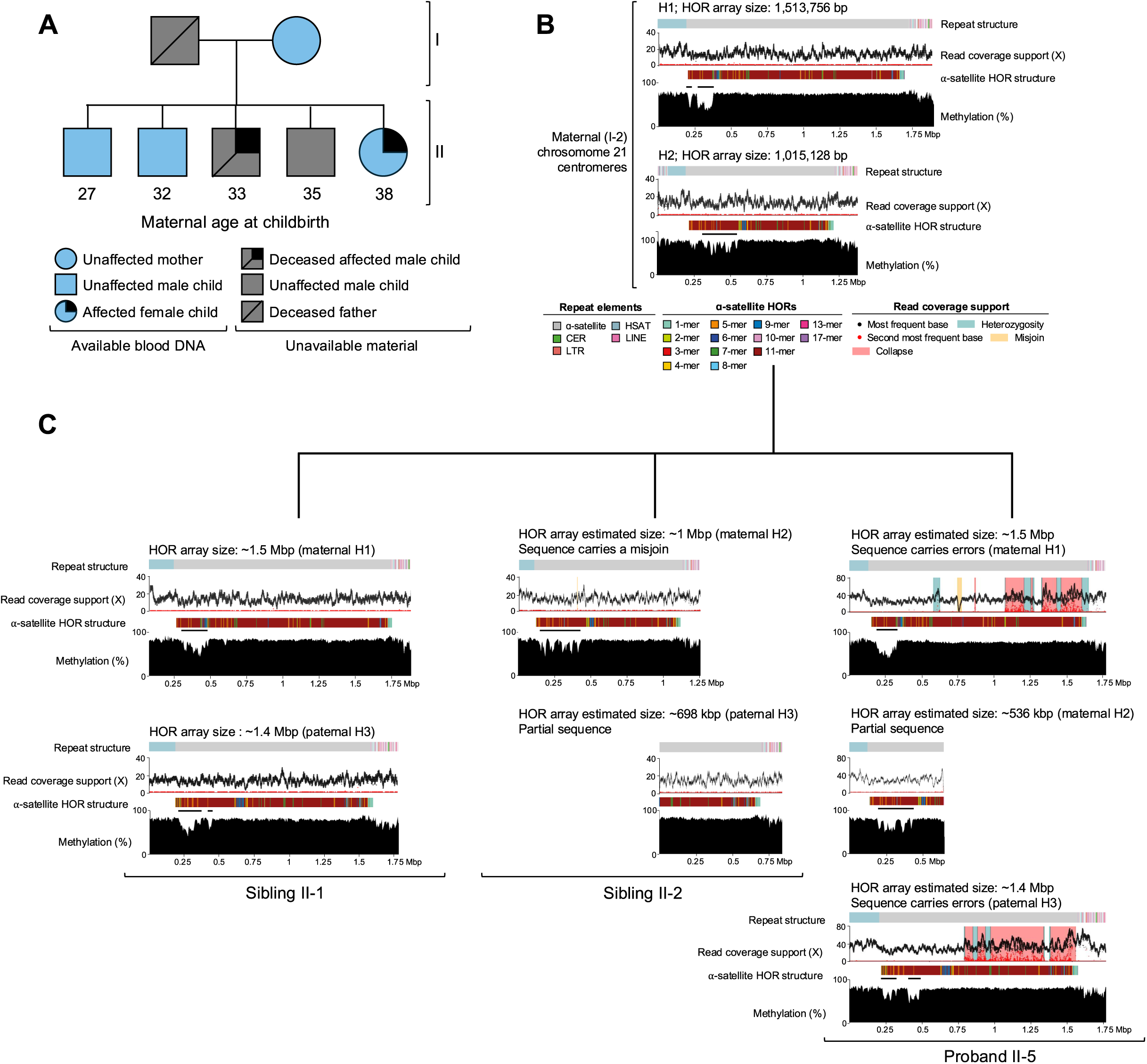
Epigenetic and centromere sequence structure of a family with recurrent T21. **A.** Family pedigree. **B.** Chromosome 21 centromeres in mother I-2. **C.** Chromosome 21 centromeres in sibling II-1, sibling II-2, and proband II-5. Repeat structure, -satellite HOR array, methylation profile, and read coverage support are shown for each centromere.

Chr21 centromeres were identified by aligning the maternal assembly contigs to the reference genome CHM13 and extracting those mapping to the chr21 centromere and q-arm as previously described (18). Both maternal chr21 centromeres are completely assembled without large-scale sequencing errors after QC (**Material and methods**) (**Fig. 1B**). We validated each assembly by aligning maternal HiFi sequence reads against each haplotype assembly showing uniform coverage and a small number of secondary small nucleotide variants (25) (**Material and methods**)—compatible with the error rate of PacBio sequencing technology. (**Fig. 1B**). This clean realignment pattern also excludes mosaicism of a third haplotype in the mother that could have predisposed her to recurrent pregnancies with T21. Both maternal chr21 centromere HOR arrays are longer than a megabase pair (Mbp) in size (H1: ~1.5 Mbp and H2: ~1 Mbp; **Table S1**) and their structure is primarily organized in canonical 11-mer units (H1: ~89%; H2: ~85%) (18) with other k-mer units present at lower frequency (**Fig. 1B**). There is no evidence of centromere size asymmetry as reported for a subset of families with young mothers with children with T21 (18).

Based on 5mC analysis of the maternal ONT data generated from blood DNA, we next defined the CDR for the two originating maternal chromosome 21 homologues using CDR-Finder (23) with standard parameters. The CDR boundary coordinates were refined using the Integrative Genomic Viewer (24) (**Material and methods**) and manual curation. We observe a typical profile for haplotype 1 (H1) with two distinct CDRs (40 and 120 kbp) and a more diffuse pattern of hypomethylation without a clear CDR on haplotype 2 (H2) over a 335 kbp region of variable methylation levels (**Fig. 1B**). We quantified the methylation levels by calculating the percentage of methylated CpGs (CpGs having more than 70% of the reads showing a methylation signature) within each CDR window (**Material and methods; Table S2**). Maternal H1 and maternal H2 show 36% and 55% methylated CpG, respectively. Thus, the maternal H2 shows an increase of 19 percentage points when compared to H1. Hereafter, we refer to the H2 CDR as a diffuse CDR (dCDR).

We next focused on the sequence and assembly of chr21 centromeres from the offspring. Biological material was available for the first two unaffected siblings, II-1 and II-2 (maternal age at childbirth: 27 and 32), and the oldest child with T21, II-5 (maternal age at childbirth: 38) (**Fig. 1A**). No biological material was available for the younger proband II-3 with T21 (maternal age at childbirth: 33) but a previous study confirmed that both affected pregnancies occurred as a result of MMIE (19) (family RDS07). A short tandem repeat analysis of the two affected individuals detected a crossover on the maternal chr21s in proband II-3 but not II-5 (**Fig. 1A**). This observation further excludes the possibility of maternal mosaicism in the population of oocytes and indicates that the two affected individuals were the result of independent nondisjunction events.

Despite generating deeper sequence coverage for proband II-5 (**Table S1**) to accommodate for the trisomy, all three centromeres are only partially assembled (**Fig. 1C**) with more errors detected when compared to the diploid mother. Consistent with the predicted MMIE (19), genome assembly analysis confirms that individual II-5 with T21 inherited the maternal H1 and H2 and a paternal haplotype (referred to as H3; **Fig. 1C**). The proband II-5 haplotypes H1 and H3 have ~1.5 Mbp (~71% correct) and ~1.4 Mbp (~53% correct) of alpha satellite assembled, respectively, while haplotype H2 is partially assembled without sequence errors for a total of ~537 kbp. All estimated HOR array sizes are reported in **Table S1** and maternally inherited centromere expected sizes are comparable with what was observed in mother II-2. The maternal transmission of both haplotypes could be reliably confirmed by aligning proband UL-ONT reads to the maternal assembly (**Fig. S1**) and via alignment of alpha satellite sequences between mother and proband, which shows sequence identity above 99% (99.4-99.9%; see **Material and Methods**). As a control, we aligned the proband II-5 paternal H3 to mother II-2 H1 and obtained a much lower sequence identity of 92.11%.

Although complete centromeres are not assembled for proband II-5, the corresponding CDR for the maternally inherited centromeres is recovered without large-scale errors. Unlike the mother, who shows two discrete CDRs on H1, the proband II-5 with T21 shows a single unique 145 kbp CDR on H1. Centromeric sequence alignment between mother and proband reveals a 3.7 kbp deletion in that region, which might explain this change. H2 has a dCDR as seen in the mother, over a region of 240 kbp and no structural variants are observed. Both haplotypes have similar methylated CpG percentages to those seen in the mother II-2 (proband H1 (35%) vs. mother’s H1 (36%); proband’s H2 with dCDR (51%) vs. mothers H2 (55%); **Table S2**). The paternal haplotype exhibits a canonical methylation profile with two CDRs (95 and 90 kbp) in close proximity (methylated CpGs: 44%) (**Fig. 1C**) (**Table S2**). Alignment of proband UL-ONT reads to the maternal assembly, other than confirming haplotype inheritance, validates the methylation profiles (**Fig. S1**).

For comparison, we also sequenced and assembled the chr21 homologs from the two unaffected siblings. Unaffected sibling II-1 inherited the maternal H1 (HOR array size: ~1.5 Mbp) with a 3.7 kbp and a 1.9 kbp deletion. Notably, the larger one is the same structural variant observed in proband II-5 H1 and may have played a role in the CDR change from two windows to one on this haplotype as well. Sibling II-1 H1 shows a slightly higher percentage of methylated CpGs (48% in sibling II-1 vs. 36% in mother) over a single 190 kbp CDR compared to the two observed in the mother (**Fig. 1C**) (**Table S2**). As before, read alignment data (**Fig. S1**) confirms the identity of the inherited chr21 centromere and its methylation profile. Centromere alpha satellite sequencing alignment also confirms 99.6% sequence identity. The II-1 sibling inherited the same paternal chr21 H3 centromere as proband II-5: a canonical HOR array structure with a size of ~1.4 Mbp and two CDR windows (160 and 28 kbp) as observed in proband II-5. Both show a similar degree of CpG methylation (47% vs. 44% in proband II-5) and, when aligned, 99.33% sequence identity.

Unaffected sibling II-2 inherited maternal H2, whose sequence is largely complete with one relatively small misjoin (~2.5 kbp). Although the precise size of the centromere HOR could not be determined, we estimate it at >1 Mbp and confirm its inheritance via alignment of UL-ONT data from sibling II-2 to the maternal genome (**Fig. S1**). Alpha satellite sequence alignment also confirms a sequence identity above 99.99%. The methylation pattern (**Fig. S1**) analysis predicts a dCDR over 295 kbp (59% methylated CpGs compared to 55% methylated CpGs in the mother; **Table S2**) This finding suggests that the diffuse methylation profile of this centromere was present in the mother since she was at least 32, a year prior to the birth of her first child (II-3) with T21 and six years before her second (II-5). The paternal chr21 centromere failed to completely assemble (assembled alpha-satellite array: ~735 kbp) and is not further assessed. However, alignment of its sequence to sibling II-1 H2 confirmed that it is the same centromere H3 (100% sequence identity).

To evaluate how common a dCDR is in the general population or if it is a phenomenon restricted to the mother II-2 chr21 H2 centromere, we assessed 16 other chromosomes where the centromeres had been sufficiently assembled in the mother (**Fig. S2**). All 16 show uniform HiFi and ONT sequence read coverage consistent with an accurate assembly and 13/16 (~81%) CDRs occur over regions without any assembler error (**Fig. S2**); 10/13 (~77%) centromeres show well-defined CDRs, while, despite not having a diffuse methylation pattern, 3/13 (~23%; chr5 H1, chr11 H2 and chr12 H2) show increased methylation peaks within the CDR windows with a methylated CpG percentage ranging from 52 to 70 (**Fig. 2A**). With these limited data and our still partial understanding of centromere methylation patterns, we cannot establish whether the dCDR on chr21 is a feature restricted to this chromosome in mother I-2.

**Figure 2.**
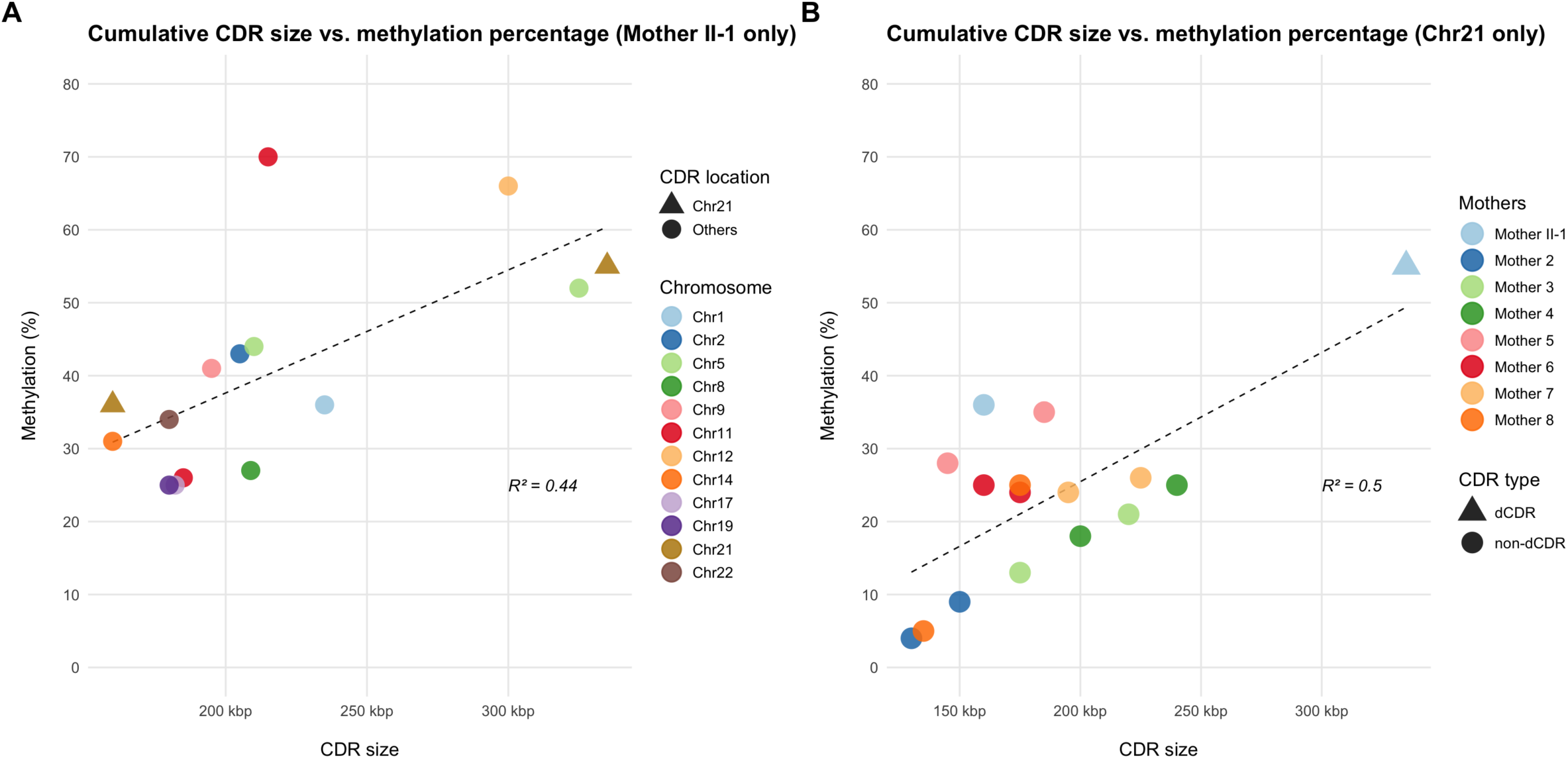
Comparative CDR analyses. **A.** Cumulative CDR sizes compared to their percentage of methylated CpGs in mother II-1 centromeres from all chromosomes. **B.** Cumulative CDR sizes compared to their percentage of methylated CpGs in mother II-1 and seven young mothers (maternal age at childbirth: ≤34) with children with T21.

To evaluate the frequency of dCDRs in other families with T21, we conducted the same analysis (**Material and methods**) on previously published chr21 centromere methylation from lymphoblastoid cell lines from seven young mothers (maternal age at childbirth: ≤34) with children with T21 (18). Among all chr21s, mother II-2 H2 represents an outlier with the largest CDR size and highest methylation levels (**Fig. 2B**). It is notable that, despite blood having generally higher methylation levels than lymphoblastoid cell lines(18), mother II-1 H1 CDR is similar in methylation and CDR size to the other mothers’ centromeres, contrarily to H2. The smallest CDRs observed in mother 2 and mother 8 are the ones present on the smallest chr21 centromeres described in females (**Fig. 2B**) (18), which are almost completely hypomethylated. In a recent population centromere study, including 108 chr21 centromere haplotypes (cencyclopedia.com) (15), no centromeres were described carrying a dCDR in lymphoblastoid cell lines. There are some population haplotypes though (e.g., HG02666-H1 and HG01457-H2; both males) that show CDRs spaced by highly methylated sequence similarly to what observed in mother II-2 chr5 H1, chr11 H2, and chr12 H2. However, they all carry deeper and more defined CDRs than mother II-2 chr21 H2 dCDR.

## DISCUSSION

The objective of this study was to assess the structure and methylation patterns of the centromere alpha satellite HOR for a family where recurrent T21 pregnancies occurred. Using long-read sequencing and one of the latest genome assembly algorithms, we reconstructed the chr21 centromeres of a mother and three of her children, one affected with T21 and two unaffected, spanning 11 years of reproductive age, a unique longitudinal perspective on centromere transmission in a family with recurrent T21 occurrences caused by MMIE (**Fig. 1**). In this family, a parental Robertsonian translocation and T21 mosaicism in maternal germ cells, either of which could explain the recurrent T21 pregnancies, are excluded. Nor did the maternal chr21 centromere homologs show evidence of size asymmetry—a feature recently observed in a subset of young mothers with children with Down syndrome (~30%) (18) and confirmed to be detrimental to female meiosis in mice oocytes (26).

We detected a transmitted change in CDRs between mother I-2 H1 and its equivalent in sibling II-1 and proband II-5. Both carry a 3.7 kbp deletion in the CDR, which likely led to the formation of a single hypomethylated region instead of the two seen in the mother. This phenomenon has been described before in a human pedigree without T21 (27), yet here we observe the same exact deletion on two haplotypes with the same consequence on the CDR. The most notable observation though is that the mother II-2 chr21 H2 lacks a clear CDR. Instead, it shows a more diffused pattern without clear dips, which we termed dCDR. Based on the other chromosome 21 centromeres observed in the family, this pattern is unusual and has a 19 percentage points difference in methylated CpGs compared to H1.

Three additional non-chr21 centromere CDRs in the mother show a high percentage of methylated CpGs, but it is unclear whether they are linked to chr21 H2 methylation increase as they also present clear and deeper hypomethylated regions (**Fig. 2A** and **Fig. S1**). The other chr21 centromeres from young mothers with children with T21 (maximum methylation percentage: 35%) show similar features to those of mother II-2 H1 with a canonical CDR. The maternal H2 instead is the only one with such a high methylation percentage and CDR size (**Fig. 2B**). Furthermore, a dCDR has not yet been described in still limited population surveys of chr21 centromeres (15), suggesting it may be a low-frequency event. Despite this evidence, contrarily to our family who was sequenced from blood DNA, all methylation data from other mothers and population samples were generated from lymphoblastoid and fibroblast cell lines, which show a lower level of centromeric methylation in a previous direct comparison (18). Also, the difference in coverage can also influence the methylation pattern observed. Therefore, our single family and the very limited availability of matching methylation data do not allow us to draw a conclusion on the frequency of dCDR in mothers with children with T21 and the general population.

The pedigree genome assembly analysis revealed that the dCDR was present on the maternal H2 since she was at least 32, a year before she had her first child with T21 and six years before her second affected child. Since sibling II-1 did not inherit that centromere and there are no earlier epigenetic data available, it is impossible to determine whether the dCDR was always present in the mother or developed with age. Despite not being fully understood, recent data are establishing the importance of CDRs and methylation profiles on centromeres and their relationship with kinetochore proteins (28,29). Recent studies confirmed an inverse relationship between methylation and the protein CENP-A, one of the main histones that defines the location and function of the kinetochore. Therefore, it is logical to hypothesize that a dCDR would have less accumulation of CENP-A and possibly, as a consequence, a weaker kinetochore that might make the chromosome more prone to nondisjunction. However, the functional importance of these methylation profiles is still unclear and other previously proposed mechanisms to explain the maternal age effects, such as reduced chiasma (5,6), cohesin deficiency (7), and additional variations of the chr21 alphoid DNA (8), could have also contributed to the T21 risk.

Based on this observation, we introduce some potential hypotheses that can be evaluated and tested in the future (**Fig. 3**). First, it is possible that women at average risk of chr21 nondisjunction carry a small fraction of oocytes at increased risk, because they carry a chr21 centromere with a dCDR (**Fig. 3B**). Alternatively, these oocytes might become at risk with age, converting their regular CDR over time to a less-defined dCDR (**Fig. 3C**). Given the depletion of oocytes during a woman’s life, the fraction of oocytes at risk might increase with the age of the woman, putting older women at an increased risk of nondisjunction. The second hypothesis is that some women might already have a higher fraction of oocytes at risk making them prone to T21 pregnancies at any age and at an increased risk with advanced age (**Fig. 3D**).

**Figure 3.**
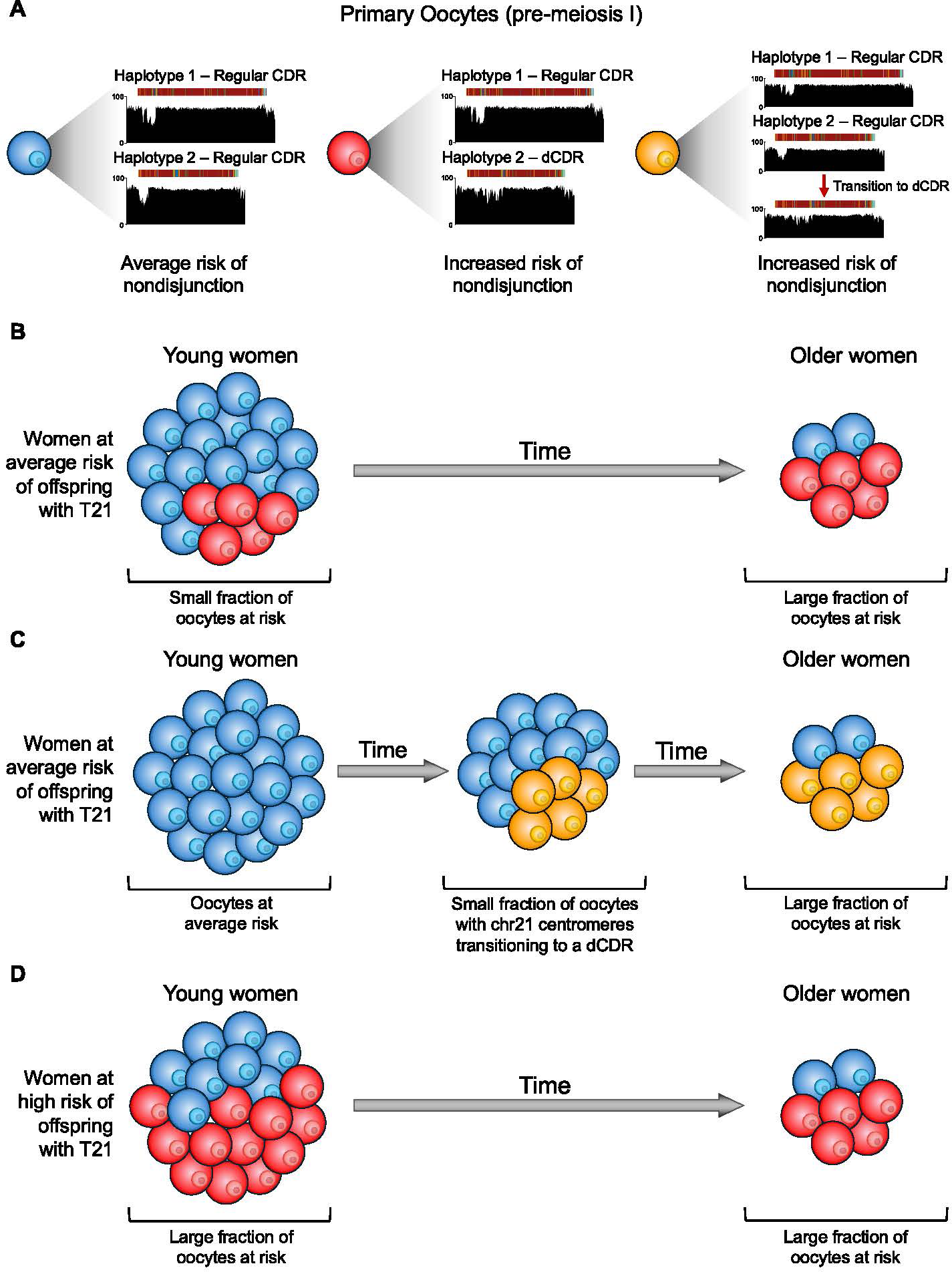
Proposed models to explain the positive correlation between maternal age and chr21 nondisjunction risk. **A.** Legend explaining the different types of oocytes. Blue has two chr21 centromeres with canonical CDRs. Red has one centromere with a canonical CDR and one centromere with a dCDR. Yellow has one centromere with a canonical CDR and one centromere whose CDR transitioned from canonical to dCDR. **B.** Hypothesis 1: Women at average risk of chr21 nondisjunction have a small fraction of oocytes at risk or **C.** a small fraction of oocytes transitioning to at-risk state over time. The percentage of oocytes at risk increases with maternal age due to progressive use of oocytes during the early reproductive years of the mother. **D.** Hypothesis 2: Some women are predisposed to chr21 nondisjunction due to a high number of oocytes at risk with dCDR. The likelihood increases over time due to the reduction of the non-dCDR oocytes available.

Several limitations should be considered when interpreting these findings. First, we only studied a small pedigree with biological samples available only for three members, which prevented the study of centromeres in the first proband and another unaffected sibling. Although recurrent T21 is exceedingly rare, other families with multiple children with T21 conceived at older age—and no parental mosaicism—should be studied at the level of genome assembly combined with epigenetic characterization to confirm that the dCDR is a shared feature of these mothers. Very young mothers with affected children should also be included, given that the meiotic segregation error risk appears to be increased in those mothers as well (<20 & ≥33) (30). Secondly, the time points of this longitudinal data are sparse (maternal age at childbirth: 27, 32, 38) and the mother’s chr21 H2 methylation profile could not be assessed when she was in her 20s. Additional families with similar features should be sequenced to compare centromeric methylation status over multiple time points. Lastly, despite having confirmed that the CDRs remain in the same position between blood and immortalized cell line DNA (18), the observed methylation levels between tissue and cell lines differ; therefore, the frequency of dCDRs should be assessed in oocytes from large population cohorts to establish whether it is a feature of mothers with children with T21 or not.

## DATA AVAILABILITY

The long-read sequencing data generated for this study were upload at the European Genome-Phenome Archive (submission ongoing).

## CODE AVAILABILITY

No custom code was generated for this study.

## ACKNOWLEDGMENTS

We thank Tonia Brown for the editing of the manuscript. Artificial intelligence was not used for the production of this manuscript.

## AUTHOR CONTRIBUTIONS

F.K.M., S.E.A., and E.E.E. conceptualized the project; S.E.A. and P.M. provided the samples; K.H. and M.A. extracted the DNA and generated the long-read sequencing data; F.K.M. performed genome assembly, sequence validation, and centromere characterization; F.K.M. managed the data submission to the European Genome-Phenome Archive. F.K.M., S.E.A., and E.E.E. drafted the manuscript. All authors read and approved the manuscript.

## ETHICAL APPROVAL

The samples in this study were obtained from a family studied in 1992 (19) (family RDS07). The family was recontacted for this analysis. After genetic counseling and explanation of the study by a medical geneticist, the family members signed an informed consent (provided by the University of Athens St. Sophia Hospital) and donated EDTA blood.

## FIGURE LEGENDS

**Figure S1.** Read coverage of UL-ONT read from each child over the maternal chromosome 21 centromeres.

**Figure S2.** Additional centromeres assembled in the mother with read coverage support and methylation profile.

## TABLE LEGENDS

**Table S1.** Summary of sequencing coverage, chromosome 21 centromere completeness, and inheritance.

**Table S2.** Quantification of methylated CpGs within chr21 CDRs for all family members.

