## Supplementary figures and images for "Sequence and epigenetic characterization of chromosome 21 centromeres in a family with recurrent Trisomy 21"

### Supplemental Figure 1

Offspring UL-ONT read alignment against the maternal assembly (mapq ≥ 5)

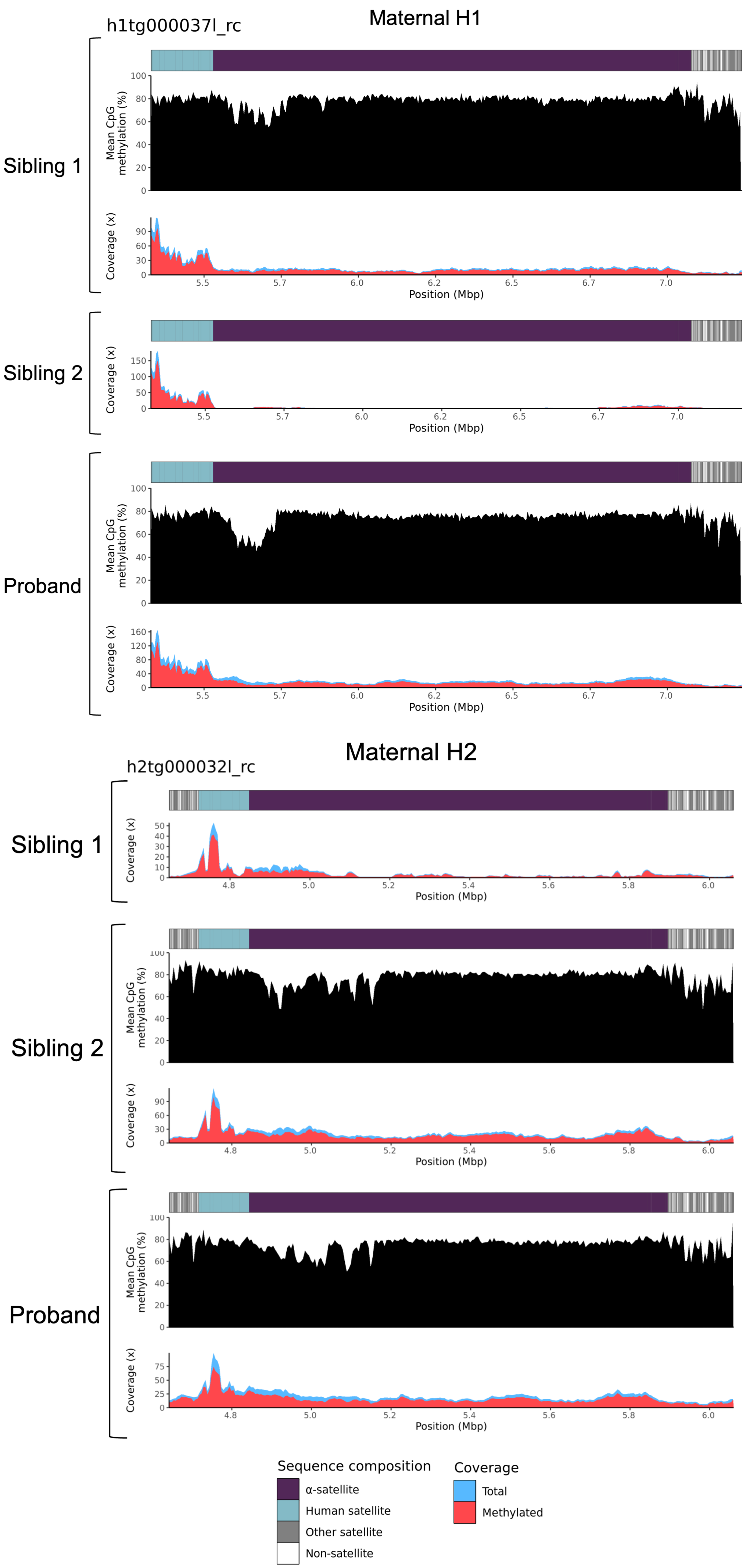

### Supplemental Figure 2

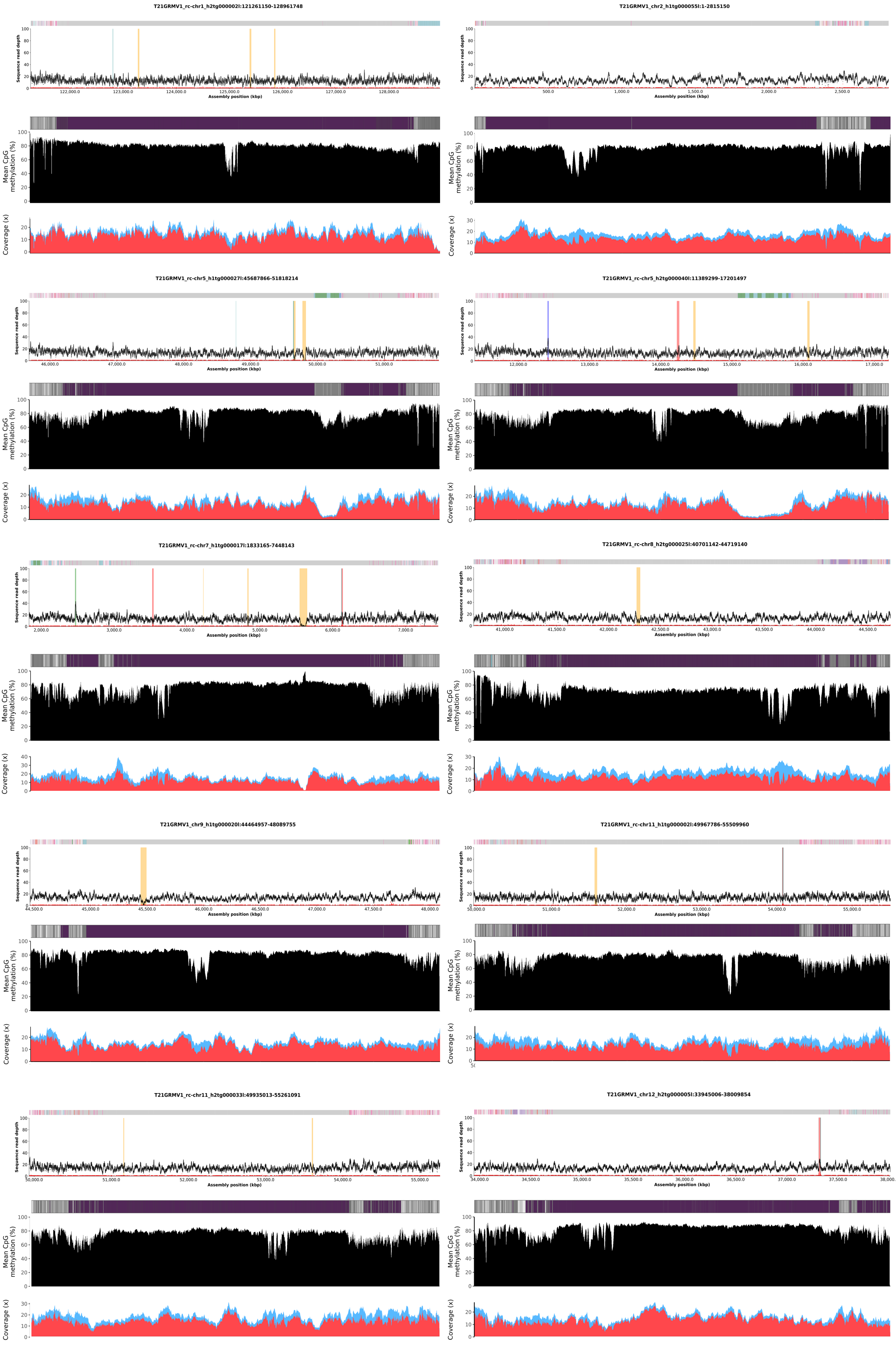

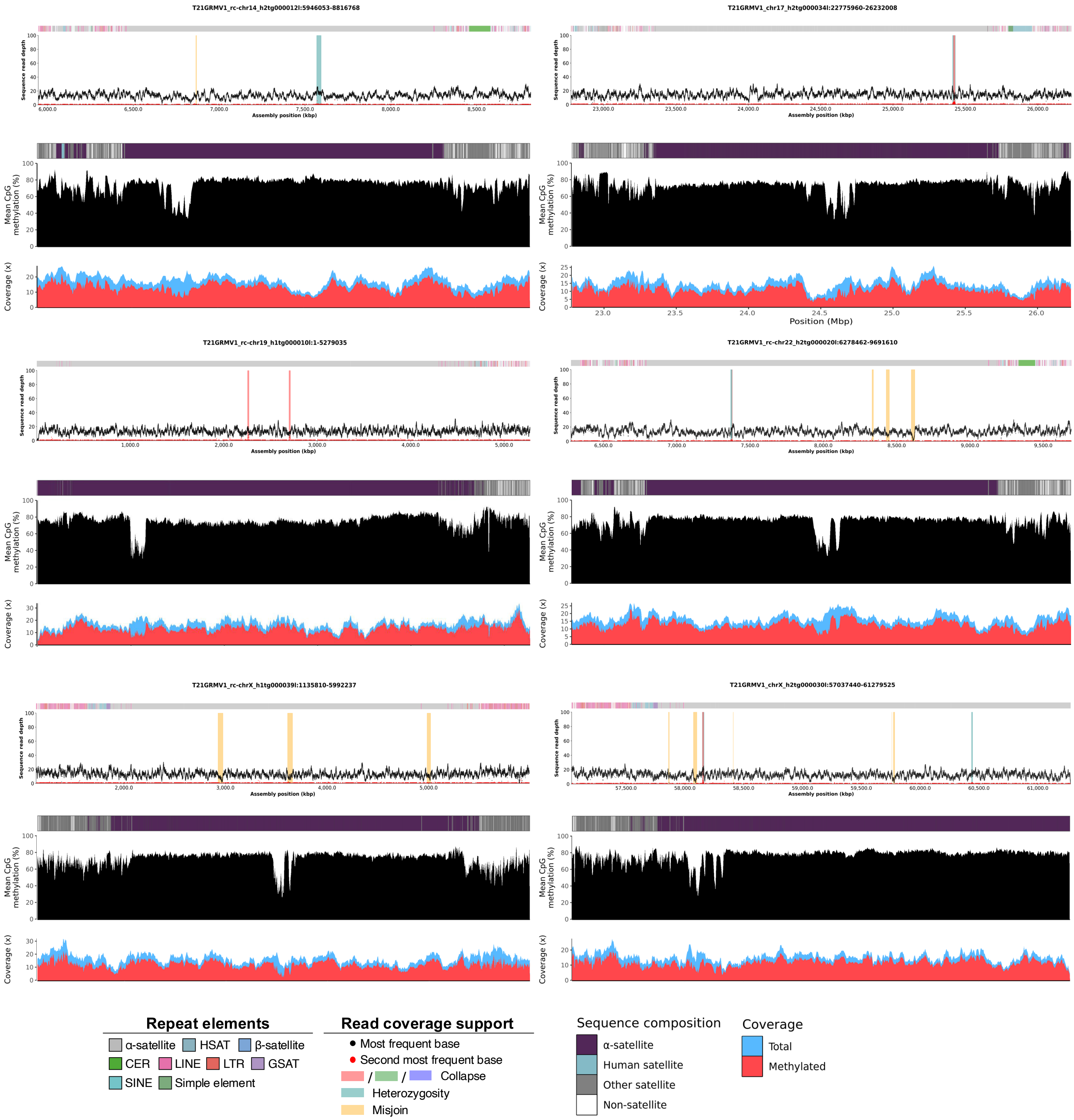
